# Identification and biochemical characterization of a novel PP2C-like Ser/Thr phosphatase in *E. coli*

**DOI:** 10.1101/303594

**Authors:** Krithika Rajagopalan, Jonathan Dworkin

## Abstract

In bacteria, signaling phosphorylation is thought to occur primarily on His and Asp residues. However, phosphoproteomic surveys in phylogenetically diverse bacteria over the past decade have identified numerous proteins that are phosphorylated on Ser and/or Thr residues. Consistently, genes encoding Ser/Thr kinases are present in many bacterial genomes such as *E. coli,* which encodes at least three Ser/Thr kinases. Since Ser/Thr phosphorylation is a stable modification, a dedicated phosphatase is necessary to allow reversible regulation. Ser/Thr phosphatases belonging to several conserved families are found in bacteria. One family of particular interest are Ser/Thr phosphatases which have extensive sequence and structural homology to eukaryotic Ser/Thr PP2C phosphatases. These proteins, called eSTPs (eukaryotic-like Ser/Thr phosphatases), have been identified in a number of bacteria, but not in *E. coli.* Here, we describe a previously unknown eSTP encoded by an *E. coli* ORF, *yegK,* and characterize its biochemical properties including its kinetics, substrate specificity and sensitivity to known phosphatase inhibitors. We investigate differences in the activity of this protein in closely related *E. coli*strains. Finally, we demonstrate that this eSTP acts to dephosphorylate a novel Ser/Thr kinase which is encoded in the same operon.

**Importance:** Regulatory protein phosphorylation is a conserved mechanism of signaling in all biological systems. Recent phosphoproteomic analyses of phylogenetically diverse bacteria including the model Gram-negative bacterium *E. coli* demonstrate that many proteins are phosphorylated on serine or threonine residues. In contrast to phosphorylation on histidine or aspartate residues, phosphorylation of serine and threonine residues is stable and requires the action of a partner Ser/Thr phosphatase to remove the modification. Although a number of Ser/Thr kinases have been reported in *E. coli*, no partner Ser/Thrphosphatases have been identified. Here, we biochemically characterize a novel Ser/Thr phosphatase that acts to dephosphorylate a Ser/Thr kinase that is encoded in the same operon.

## Introduction

Reversible protein phosphorylation is an important regulatory mechanism in eukaryotes and prokaryotes (1). In eukaryotes, signaling phosphorylation typically occurs on serine, threonine or tyrosine residues and is mediated by the combined action of kinases and phosphatases. In prokaryotes, signaling phosphorylation has been thought to occur largely on histidine and aspartate residues mediated by histidine kinases of two-component systems (2). However, mass spectrometry based-phosphoproteomic analyses over past decade have identified numerous Ser/Thr/Tyr phosphorylated proteins in many bacteria (3–5), including *Escherichia coli* (6–9). Some of these phosphoproteins and the specific phosphosites are conserved in divergent species (7) suggesting that this regulation may be physiologically relevant.

Ser/Thr kinases from phylogenetically diverse bacteria have been described (5). For example, in *E. coli*, YeaG plays a role in nitrogen starvation (10), YihE is involved in the Cpx stress response (11) and cell death pathways (12) and HipA regulates bacterial persister formation by phosphorylating a tRNA synthetase (13, 14). However, both the authentic *in vivo* substrates of these kinases and/or their proximal activating stimuli are largely uncharacterized, complicating efforts to understand their precise physiological role.

Phosphorylation on serine or threonine residues is more stable than phosphorylation on histidine or aspartate residues and is subject to additional regulation by Ser/Thr phosphatases. Analysis of phylogenetically diverse bacterial genomes revealed the presence of genes encoding proteins (15–17) which bear significant resemblance to eukaryotic Ser/Thr PP2C phosphatases (17, 18) hence they are referred to as eukaryotic-like Ser/Thr phosphatases (eSTPs). Some of these proteins have been characterized biochemically and structurally, with these studies confirming their general similarity to their eukaryotic counterparts. Eukaryotic Ser/Thr protein phosphatases are divided into two classes, either phosphoprotein phosphatases (PPP) or metal dependent protein phosphatases (PPM), according to structure, presence of signature motifs, metal ion dependence and sensitivity to inhibitors (19). PPM phosphatases require Mg^2+^/Mn^2+^ to mediate dephosphorylation of phospho-serine or phospho-threonine residues. A well-studied member of the PPM phosphatase family is human protein phosphatase 2C (PP2C) (20) which bears a striking resemblance to bacterial PPM phosphatases. PPM/PP2C phosphatases are characterized by the presence of 11 signature motifs with 8 absolutely conserved residues (17, 19).

While several bacterial PPM-type phosphatases have been biochemically characterized (21–28), PPM-type phosphatases have not been identified in *E. coli* despite strong evidence of Ser/Thr phosphorylation. Here, we characterize a previously undescribed ORF, *yegK*, that is present in both *E. coli* B and K strains. This ORF encodes a ~28 kDa protein which bears sequence similarity to PP2C-type phosphatases. We designate this gene *pphC* and its protein product as PphC. Recombinant PphC was purified and its enzymatic properties characterized. Despite some differences in sequence conservation as compared to other bacterial eSTPs, PphC resembles PP2C-type phosphatases in various biochemical assays. Furthermore, we show that PphC dephosphorylates autophosphorylated YegI (a previously undescribed Ser/Thr kinase) which is encoded in the same operon as *pphC.* To our knowledge, this is the first report of the identification and biochemical characterization of an *E. coli* PP2C-like phosphatase.

## Results

### YegK is an atypical PP2C-like phosphatase

To identify PP2C-like phosphatases in *E. coli*, we performed a homology-based BLAST search using *Bacillus subtilis* PrpC, a well-characterized PP2C-like phosphatase (26), as the query. This analysis revealed a previously uncharacterized 762bp ORF, *yegK*, which encodes a putative 253 amino acid polypeptide. Sequence analysis and domain prediction of YegK revealed that amino acids 11–232 have homology to a PP2C domain (Fig 2A). PP2C phosphatases include 11 conserved signature motifs (17, 29). Multiple sequence alignment of YegK with known PP2C phosphatases from other Gram-negative bacteria shows presence of these 11 motifs but with low overall sequence homology. However, unlike other bacterial PP2C homologs (22, 23, 25, 28, 30), YegK contains only six of the eight absolutely conserved residues that are involved in metal binding, coordination and catalysis (Fig. 1A). In particular, the amino acid sequence alignment clearly shows that YegK lacks the conserved glycine residue in motif VI and the aspartic acid residue in motif VIII (Fig. 1A).

**Figure 1:**
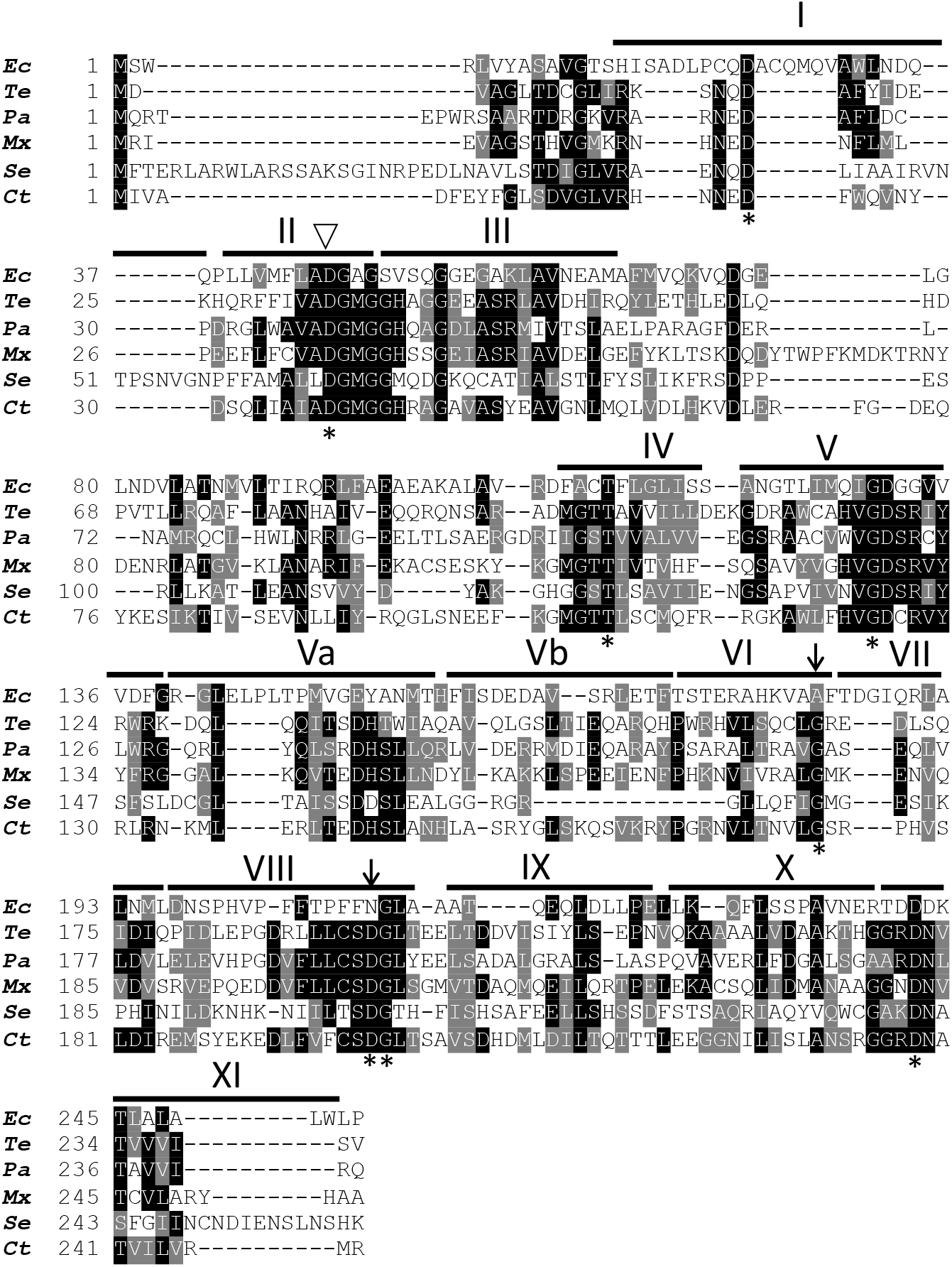
YegK is an atypical PP2C-like phosphatase. Amino acid sequence alignment of YegK with bacterial PP2C-like phosphatases. YegK *(Ec: Escherichia coli)* was aligned to PP2C homologs from Gram-negative bacteria using T-coffee (51) and Boxshade. Identical residues are shaded in black and similar residues are shaded in gray. The eight absolutely conserved residues found in bacterial PP2Cs are depicted with asterisks. The conserved residues absent in YegK are indicated by arrows. Signature motifs seen in most bacterial PP2Cs are denoted as roman numerals based on Shi et al. (17). The aspartic acid residue involved in metal binding is depicted with open triangle. *Te: Thermosynecococcus elongatus* tPphA; *Pa: Pseudomonas aeruginosa* Stp1; *Mx: Myxococcus xanthus* Pph1; *Se: Salmonella enterica* serovar Typhi PrpZ (aa1-260); *Ct: Chlamydia trachomatis* CTL0511 (Cpp1).

A predicted structure of YegK using Phyre2 algorithm (Fig. S1) closely resembles the published structure of the bacterial PP2C-like phosphatase PphA from *Thermosynechococcus elongatus* (32) consisting of ß-sheets and a catalytic core in the center surrounded by exterior alpha helices. In addition, *yegK* is located immediately upstream of *yegl*, an ORF which encodes a protein with homology to eukaryotic-like Ser/Thr kinases, consistent with the observation that bacterial Ser/Thr kinases and phosphatases are often located in operons (5). Taken together, these observations suggest that, despite the absence of two highly conserved residues, *yegK* likely encodes a PP2C-like phosphatase.

### Biochemical characterization of YegK

To demonstrate that *yegK* encodes a functional protein phosphatase, the 762bp fragment was cloned in frame with a N-terminal six-histidine tag into the pBAD24 vector (33). The plasmid was transformed into *E. coli* C43 (DE3) and following protein expression, the cell lysate was subjected to affinity purification using Ni^2+^-NTA resin and subsequent analysis by SDS-PAGE. The protein migrated at an apparent molecular mass of ~25kDa, similar to the calculated molecular mass of 28.18kDa (Fig S2). The phosphatase activity of purified YegK was determined using an absorbance-based assay which measures the hydrolysis of p-nitrophenyl phosphate to p-nitrophenol. Formation of p-nitrophenol detected at an absorbance of 405nm is directly proportional to the phosphatase activity which is expressed as nmol of pNP formed/μg protein. YegK displayed a time-dependent increase in phosphatase activity (Fig. 2A) consistent with the alignment (Fig. 1), that it is an active protein phosphatase. We have therefore renamed YegK as PphC (Protein phosphatase C) following the nomenclature of two previously characterized *E. coli* protein phosphatases PphA and PphB (34) and the bacterial PP2C-like phosphatase PphA from *T. elongatus (32).*

**Figure 2:**
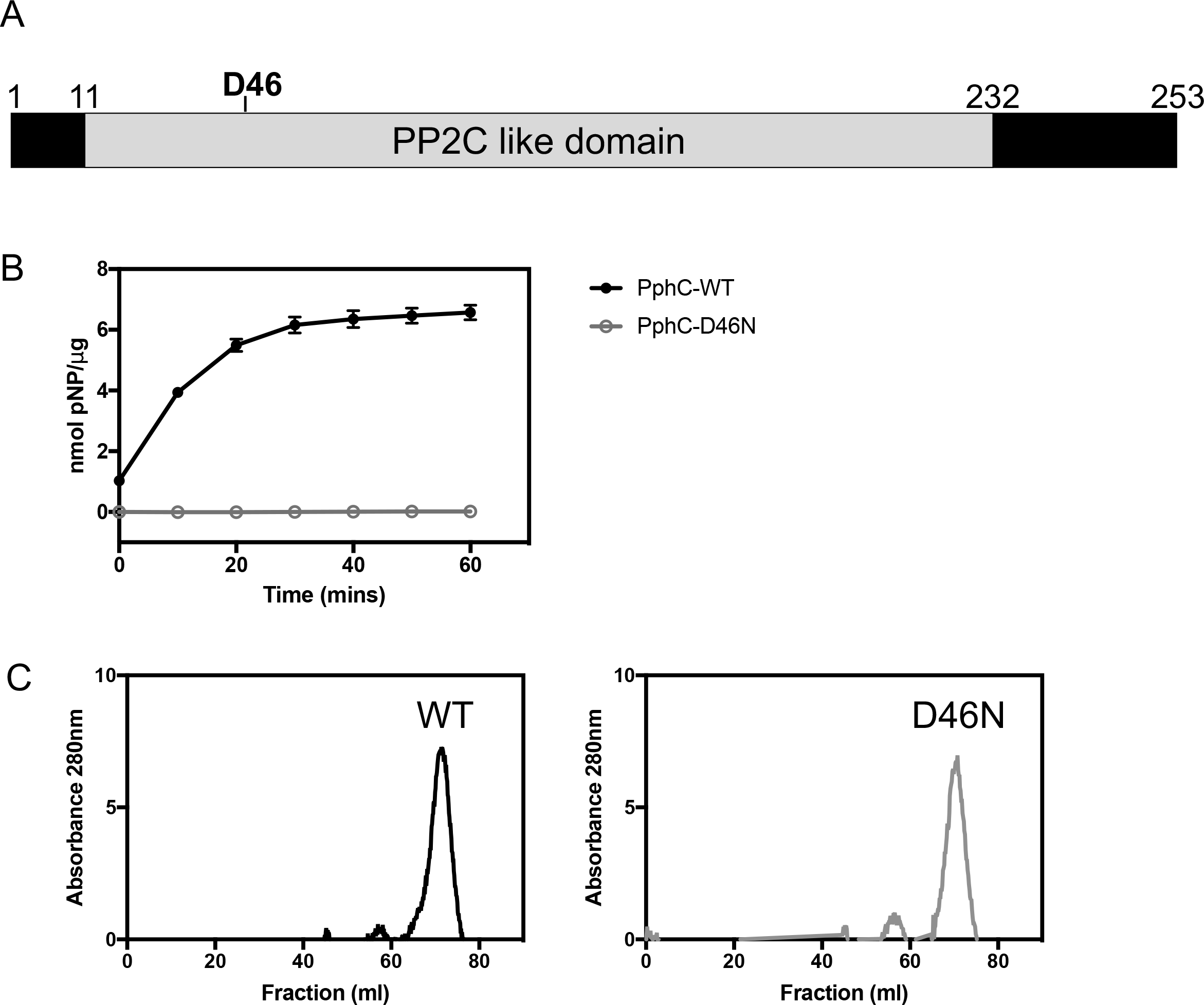
YegK is an active phosphatase. (A) Domain architecture of YegK. Metal binding site D46 is highlighted in black. (B) Assessment of phosphatase activity of YegK using pNPP as substrate. Reactions were performed at 37 °C for 60 mins with 350 nm of phosphatase (WT/D46N mutant), 5mM pNPP substrate in phosphatase assay buffer (20mM Tris pH 8.0 and 5mM MnCl2). (C) Size exclusion chromatography profile of WT and D46N YegK. His6-tagged protein (10nmol) was loaded on a Superdex 75 size exclusion column and eluted in 20mM Tris pH 8.0, 50mM NaCl, 1mM DTT, 1mM MnCl2.

To further confirm the bioinformatic identification of PphC as a PP2C-like phosphatase, the aspartic acid residue (D46) in motif II was mutated to asparagine. A similar approach was used in the analysis of the PP2C-like phosphatase Cpp1 from *Chlamydia trachomatis* (22). The mutant protein (PphC-D46N) was purified as above and phosphatase activity was compared to wild type PphC. PphC-D46N displays no hydrolysis of pNPP suggesting that the aspartic acid residue is essential for catalytic activity (Fig. 2B). To ensure that the loss of activity of PphC-D46N was not a consequence of improper folding, PphC and PphC-D46N were subjected to size exclusion chromatography. The gel filtration elution profile shows that PphC-D46N eluted as a single peak and at the same retention volume as PphC indicating that the loss of phosphatase activity observed with the PphC-D46N mutant is most likely due to a loss of catalytic activity (Fig. 2C).

PP2C phosphatases belong to the PPM family of metal dependent Ser/Thr phosphatases that require either Mg^2+^ or Mn^2+^ for activity (19, 29). The requirement for a divalent cation for PphC phosphatase activity was assessed by measuring hydrolysis of pNPP in the presence of either MgCl2/MnCl2/ CaCl2/ ZnCl2, NiCl2. Since pNPP hydrolysis was only observed in the presence of MnCl2 but not MgCl2,CaCl2, ZnCl2, NiCl2, PphC is Mn^2+^ dependent phosphatase (Fig. 3A; Fig. S3). This result is consistent with the requirement for Mn^2+^ ion for previously characterized bacterial PP2Cs (21–25, 27). The concentration dependence of PphC phosphatase activity on Mn^2+^ was measured and the optimal MnCl2 concentration was determined to be 5mM (Fig. 3B).

**Figure 3:**
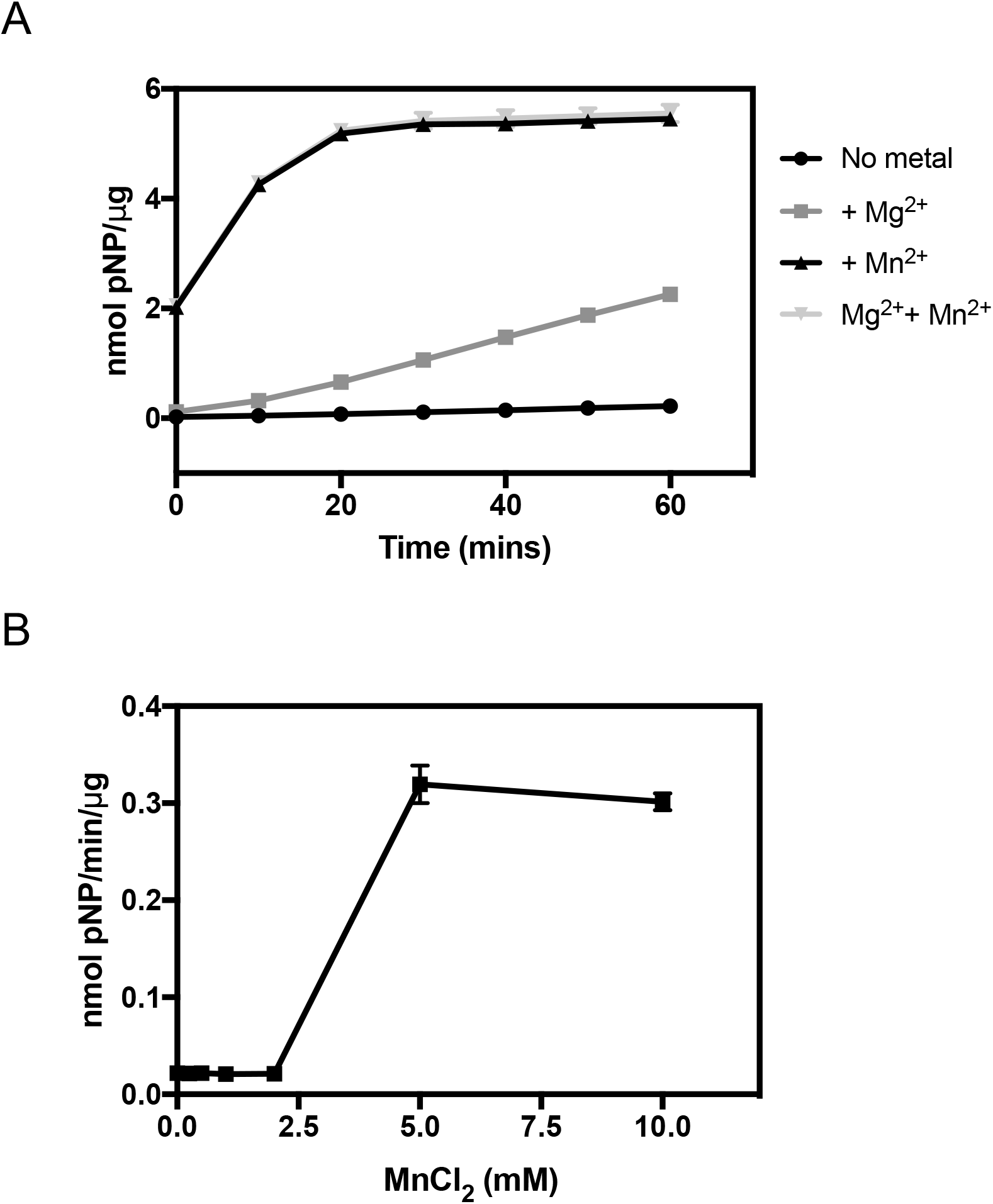
PphC (YegK) is a Mn^2+^-dependent PP2C-like phosphatase. (A) Metal dependency of PphC phosphatase was tested using pNPP as substrate. Reactions were carried out at 37 °C for 60 min in buffer containing 350 nM phosphatase, either 5mM MgCl2 or 5mM MnCl2, or both, with 5mM pNPP substrate. (B) Effect of Mn^2+^ concentration on YegK catalytic activity using pNPP as a substrate. Reactions were carried out at 37 ºC for 30 min in assay buffer containing 350 nM phosphatase, 5mM pNPP substrate and MnCl2.

The effect of different classes of protein phosphatase inhibitors on PphC phosphatase activity was tested using pNPP as a substrate. PphC activity dramatically decreased in the presence of general protein phosphatase inhibitor sodium phosphate and was slightly affected by sodium fluoride (~30% decrease at 100mM). Okadaic acid, a known inhibitor of PP2A and PP2B family of phosphatases (35), did not inhibit PphC activity. PphC activity was also unaffected by sodium orthovanadate, a known tyrosine phosphatase inhibitor. Aurin tricarboxylic acid and 5,5’ Methylene disalycilic acid which were previously reported to inhibit *Staphylococcus aureus* Stp1 (36, 37) had little effect on PphC phosphatase activity. Similarly, sanguinarine chloride an inhibitor of Human PP2Ca (38) did not affect phosphatase activity. However, a ~60% decrease in activity was detected in the presence of bivalent metal chelator EDTA confirming that PphC is a metal dependent phosphatase. Together, these data are consistent with the characterization of PphC as a PP2C-like phosphatase.

### PphC phosphatase activity is different in closely related *E. coli* strains

We identified striking differences in the genetic architecture surrounding the *pphC (yegK)* locus in *E. coli* K and B strains. Specifically, in the *E. coli* B strain REL606, *pphC (yegK)* is immediately upstream *of yegI*, an ORF encoding a putative Ser/Thr kinase, whereas in the *E. coli* K strain MG1655, a putative ORF, *yegJ*, is located between the *pphC (yegK)* and *yegI* genes facing the opposite direction (Fig. 4A). This different genomic organization is conserved in other K and B strains, suggesting that it pre-dates the divergence of these lineages (39). A reasonable prediction is that *pphC* and/or *yegI* expression would be affected by the presence of the divergently oriented *yegJ*, but as we have not been able to identify conditions under which we can robustly detect *pphC* expression, it has not been possible to evaluate this prediction.

**Figure 4:**
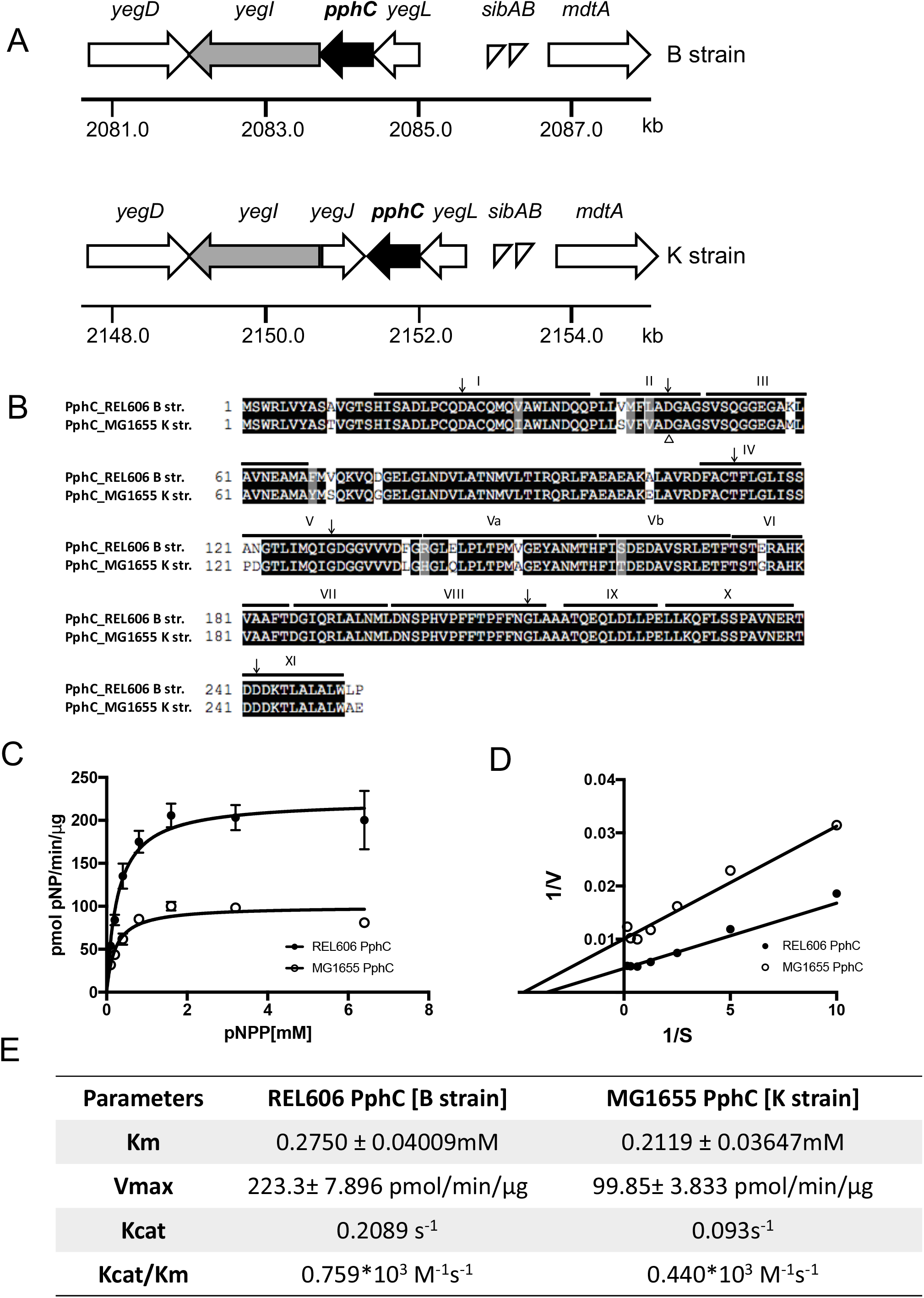
PphC phosphatase activity differs in closely related *E. coli* strains. (A) Genetic map of *yegK/yegI* operon from *E. coli* B strain REL606 and K strain MG1655: Thick arrows denote ORFs. The genes *yegL, yegK* and *yegI* encode a von Willebrand factor type A, a PP2C-like phosphatase and an eukaryotic-like Ser-Thr kinase respectively. The genes *mdtA, mdtB, mdtC* encode subunits of a multidrug efflux pump and *yegD* encodes an actin family protein. The *yegJ* gene encodes a protein of unknown function. (B) Amino acid sequence alignment of PphC from *E. coli* B strain REL606 and K strain MG1655. Identical residues are shaded in black and similar residues are shaded in gray. Signature motifs seen in bacterial PP2C phosphatases are depicted as Roman numerals based on (17). The aspartic acid residue involved in metal binding is conserved and indicated by open triangle. The conserved residues found in bacterial PP2Cs are depicted with arrows. (C and D) Enzyme kinetics of PphC. Substrate dependent activity was assessed using different concentrations of pNPP (0.1–6.4mM) in assay buffer containing 350nm phosphatase and 5mM MnCl2. Data were fitted to a Michaelis-Menten curve (C) and a Lineweaver-Burk plot (D). The Lineweaver-Burk plot was used to determine Km, Vmax and Kcat values (E).

In addition to this difference in the genetic architecture around the *pphC* locus, multiple sequence alignment of *pphC* from a K strain (MG1655) and a B strain (REL606) revealed that only ~92% of the PphC sequence is conserved (Fig. 4B). This is in contrast to the extremely high degree of sequence identity typically observed for the *E. coli* proteome: more than half the proteins in MG1655 have 100% sequence identity with the corresponding protein in REL606 (39). To determine whether these differences in amino acid sequence affect phosphatase activity, the gene product of *yegK* from *E. coli* MG1655 was over-expressed and purified by the same method used to purify REL606 PphC. The enzyme kinetics of the two proteins was compared using the pNPP assay (Fig. 4C). Phosphatase activity (pmol of pNP/min/μg) was determined with increasing concentrations of pNPP and Km and Vmax values were calculated (Fig. 4D). Rel606 PphC is more active than MG1655 PphC (Fig. 4E) since its Km and Vmax are lower, indicating that it reaches a maximum velocity at a lower substrate concentration. Similarly, the Kcat values were calculated to be 0.2089 s^−1^ and 0.093s^−1^ for REL606 and MG1655 PphC, respectively. Previously reported kinetic values for known bacterial PP2Cs range from 0.35mM to 5.7mM pNPP for Km and 0.1 to 7.4 s^−1^ for Kcat (22–24, 26, 27, 31, 40–43), indicating that PphC has relatively low phosphatase activity *in vitro* as compared to previously characterized bacterial PP2C-like phosphatases.

### Substrate specificity of PphC

The results of the pNPP assay suggested that PphC has phosphatase activity. Therefore, we investigated whether PphC is capable of dephosphorylating a protein substrate. ß- casein is phosphorylated on five serine residues at the N-terminus (44) and was used as a model protein in our assay. Using Mn^2+^-Phos-tag/SDS-PAGE to monitor phosphorylation state (45), untreated ß-casein migrated at an apparent molecular mass of 30kDa (Fig. 5A, lane 1), but ß-casein that had been pre-incubated with PphC exhibited a mobility shift (Fig. 5A, lane 2). Since such a change is indicative of a loss in phosphorylation, PphC likely dephosphorylated the serine residues of ß-casein. Further, this mobility shift was not detected following incubation of ß-casein with catalytic mutant PphC-D46N (Fig. 5A, lane 4). These results indicate that PphC is capable of acting as a serine phosphatase.

**Figure 5:**
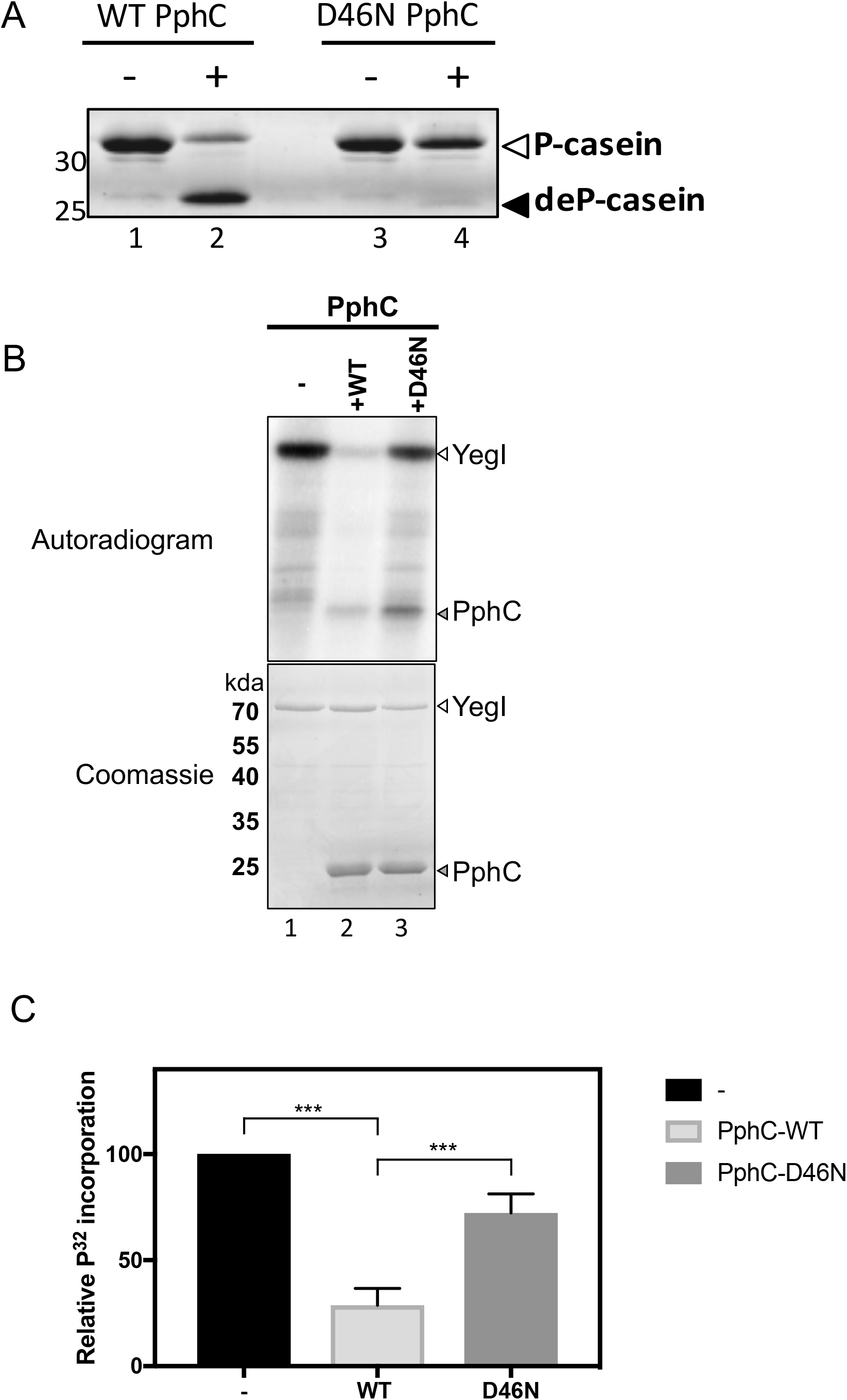
Substrate specificity of PphC (A) Effect of PphC on phospho ß-casein as a substrate. Dephosphorylation reactions were carried out at 37 ºC for 60 min in buffer containing 50mM Tris pH 8, 10mM MnCl2, 4μg phosphorylated ß-casein, 1.5μg (9μM) phosphatase. (B) Effect of PphC on YegI kinase. Dephosphorylation reactions were carried out at 37 ºC with 2μM of phosphorylated YegI and 4μM PphC in reaction buffer as described. Reactions were stopped at t=60mins and run on 12% SDS-PAGE followed by autoradiography. Molecular weights are denoted as kDa. (C) % relative ^32^P incorporation was calculated by densitometry analysis using FIJI software. Data represents the mean +/- SE of five independent experiments. Statistical analysis used unpaired T-test (***P value<0.0001).

The substrate specificity of PphC was further examined using commercially available phosphopeptides. Previously characterized PP2Cs have demonstrated preferential specificity to phosphoserine/threonine peptides over phosphotyrosine peptides (22, 23, 30, 40). Phosphatase assays were performed with phosphoserine RRA(pS)VA, phosphothreonine KR(pT)IRR and phosphotyrosine RRLIEDAE(pY)AARG peptide substrates (Fig. S4). PphC released two-fold more phosphate in the presence of the phosphotyrosine peptide as compared to the phosphothreonine peptide and it had no effect on the phosphoserine peptide. To confirm that the phosphopeptides are capable of being dephosphorylated, we used a known PP2C-like phosphatase *(B. subtilis* PrpC) as a positive control. As expected, PrpC dephosphorylated both serine and threonine residues and had minimal activity to tyrosine. Thus, in comparison to PrpC, PphC had overall minimal activity on phosphopeptides suggesting that they may not be an ideal substrate for PphC.

### Identification of a PphC substrate

Bacterial PP2C-like phosphatases are often present in the same operon as a Ser/Thr kinase (5) and the kinase is often itself a substrate of the phosphatase (25–27, 46). As noted above, *pphC(yegK)* is located immediately adjacent to *yegI*, a 1947bp ORF encoding a 648 amino acid protein with homology to eukaryotic-like Ser/Thr kinases (Fig. 4A). (5). Thus, to examine if YegI could serve as substrate for PphC, autophosphorylated YegI kinase (Fig. 5B, lane 1) was incubated in the presence of wild type or D46N mutant PphC and assayed for loss of phosphorylation. While wild type PphC dephosphorylates YegI kinase, as indicated by the loss of the radiolabeled band (Fig. 5B, lane 2; 5C), this effect is not seen following incubation with the catalytic mutant PphC-D46N (Fig. 5B, lane 3; 5C).

In these assays, a phosphorylation product of ~25kDa corresponding to PphC-D46N was observed (Fig. 5B, lane 3), suggesting that YegI could also phosphorylate PphC. However, we have been unable to demonstrate any effect of this modification on PphC activity using either the pNPP or the ß-casein dephosphorylation assays (data not shown). Interestingly, while PphC-D46N is phosphorylated, phosphorylation of wild type PphC is not observed (Fig. 5B, lane 2), suggesting that PphC can dephosphorylate itself.

### Discussion

Mass spectrometry-based phosphoproteomic analysis has revealed that many Ser/Thr/Tyr residues undergo phosphorylation in phylogenetically diverse species. In most cases, the kinases and phosphatases responsible for these modifications have not been identified, although the presence of homologs of eukaryotic Ser/Thr kinases and phosphatases in most (if not all) bacterial species suggest that they could play a role. These so-called eukaryotic-like Ser/Thr kinases and their partner PP2C-like Ser/Thr phosphatases have extensive structural and biochemical similarity with their eukaryotic counterparts (5). In *E. coli*, extensive Ser/Thr phosphorylation has been observed, with several studies reporting >75 proteins (6–9). However, the kinases and phosphatases responsible for making or removing these modifications are not known. Here, we describe biochemical analysis of PphC, a protein product of a previously undescribed *E. coli* ORF, *yegK*, that encodes the first reported PP2C-like Ser/Thr phosphatase in *E. coli.*

PphC contains all of the 11 conserved motifs present in PPM/PP2C phosphatases (16, 17,19, 29). However, unlike bacterial PP2C-like phosphatases from other Gram-negative bacteria, PphC has only 6 out of the 8 absolutely conserved residues involved in metal co-ordination and catalysis (32, 43, 47). Specifically, PphC lacks a conserved glycine residue in motif VI and a conserved aspartate residue in motif VIII (Fig. 1A). The aspartate residue in motif VIII is important for metal ion co-ordination in bacterial PP2C-like phosphatases (32, 43, 47). Despite these differences in amino acid sequence, PphC was able to effectively hydrolyze the chromogenic substrate pNPP, suggesting that the requirement of all 8 residues as a criterion for assessing the likelihood that an ORF encodes a PP2C-like phosphatase may be too stringent.

The regulation of PP2C-like phosphatases has been studied in a number of contexts including the *Mycobacterium tuberculosis* PstP PP2C-like phosphatase whose activity is stimulated by phosphorylation on multiple Thr residues by two eukaryotic-like Ser/Thr kinases (48). Although we observed that PphC is phosphorylated by its partner Ser/Thr kinase YegI (Fig. 5B), we were unable to detect any effect on PphC activity. Another regulatory mechanism occurs in the *B. subtilis* SpoIIE protein, a PP2C-like phosphatase that plays an essential role during sporulation. A single residue in SpoIIE (Val-697) mediates an alpha-helical switch that changes the coordination of a metal ion in the active site and thereby activates the phosphatase (49). However, since this residue is not conserved in PphC, this regulatory mechanism is probably not operating in the context of PhpC function.

Bacterial PP2C-like phosphatases are known to dephosphorylate pSer/pThr containing peptides (22–24). PphC was initially identified using a homology search using *B. subtilis* PrpC but unlike PrpC, PphC failed to dephosphorylate phosphopeptides and displayed no preference for pSer/pThr/pTyr peptides (Supp. Fig. 4). This lack of activity against phosphopeptides is in contrast to the ability of PphC to dephosphorylate the pSer containing protein substrate ß-casein. A possible explanation could be that PphC may require additional residues for substrate recognition and/or binding that are not present in the phosphopeptides. However, this is not likely to be a sufficient explanation as ß-casein is a generic substrate that would not be expected to contain specificity determinants for PphC. Alternatively, this result suggests that there may be limits to using phosphopeptide dephosphorylation as an accurate assay of PP2C-like phosphatase function.

Bacterial eukaryotic-like Ser/Thr kinases and PP2C-like phosphatases are often encoded in the same operon and in many cases the kinase is a substrate for the phosphatase (21, 25–27). Similarly, in the *E. coli* B strain REL606, the ORF *yegI* encodes a putative eukaryotic-like Ser/Thr kinase and is located immediately downstream of *pphC.* Our data demonstrate that YegI is a substrate of PphC (Fig. 5C). However, in *E. coli* K strain MG1655, there is a putative intervening ORF, *yegJ*, located between the *yegI* and *pphC* genes and that is transcribed divergently (Fig.4A), suggesting potential regulatory differences between the two strains. We have been unable to observe transcription of the *pphC* locus under any conditions, so we do not know if *yegJ* has an effect on expression of this locus. Interestingly, mutations in *yegI*, the gene encoding the partner kinase of PphC, repeatedly emerge in long-term evolution experiments (50), suggesting that this kinase/phosphatase pair may have significant fitness effects. Consistently, the presence of the potentially disruptive *yegJ* locus in the K lineage may be also reflect these effects. However, at present, in the absence of a deeper understanding of the physiological role of YegI/PphC pair, these effects remain mysterious.

In addition to the differences in the genome organization around *pphC* in the *E. coli* B and K lineages, the amino acid sequence of PphC differs between the two strains, with several non-conservative substitutions. This observation is intriguing given that half of the proteome is identical between these two strains (39). Although these differences are not in residues that are absolutely conserved among bacterial PP2C-like phosphatases (32, 43, 47), they may have functional consequences since the two proteins exhibited modest differences in enzyme kinetics (Fig 4C-E). Future work will be aimed at identifying the impact of specific substitutions in the residues that differ on the relative activity of the different PphC alleles.

Finally, PP2C phosphatases play important roles in cellular regulation in more complex systems including mammals (19). One issue in investigating PP2C function in these *in vivo* contexts is that specific chemical inhibitors do not exist. Thus, the bacterial homologs such as PphC may be useful in identifying cell-permeable inhibitors, both because of the technical tractability of the organism as well as the presence of only a single PP2C-phosphatase gene in the genome.

In summary, our study provides the first evidence for the existence of a PP2C-like phosphatase in *E. coli.* Despite some differences in sequence conservation as compared to other PP2Cs, PphC is an active PP2C-like phosphatase, albeit with lower Km and Vmax values than other bacterial PP2Cs. Future studies will be required to identify physiological substrates of PphC and to ascertain its physiological role *in vivo.* We expect that further characterization of PphC’s partner kinase, YegI, will greatly facilitate these efforts.

### Experimental Procedures

#### Bacterial strains and growth conditions

*E. coli* DH5a cells were used for regular cloning and C43(DE3) and LOBSTR (BL21-DE3) strains were used for expression of recombinant proteins. *E. coli* cells were grown in LB Lennox broth supplemented with ampicillin (100μg/ml) at 37 °C with shaking (220 rpm) unless otherwise specified. Details of strains, plasmids and primers used in the study are described in Supplementary Tables S1, S2, and S3, respectively.

#### Cloning and expression of YegK(PphC)

Genomic DNA from *E. coli* REL606 and MG1655 was isolated using a Wizard Genomic DNA purification kit (Promega) following manufacturer’s instructions. The *E. coli yegK (pphC)* gene was PCR amplified using Phusion polymerase (Thermo Scientific) from REL606 genomic DNA using primers (KR38/KR39). Sequence for an N-terminal His6 tag was included in the primer. The PCR product was digested with NcoI/PstI and ligated into similarly digested pBAD24 plasmid backbone. Ligation products were transformed into DH5a cells and selected on LB ampicillin plates. The resulting plasmid generated an N-terminal His6-tagged YegK (PphC) fusion protein.

For protein expression, all plasmids were transformed in C43DE3 cells and plated on LB ampicillin (100μg/ml) agar plates. Single colonies were inoculated into 3ml LB supplemented with ampicillin (100μg/ml) for overnight cultures. The next day, 400ml cultures were diluted 1:250 and grown to an OD600 of 0.6–0.8. Recombinant protein was induced by addition of arabinose (0.2% w/v) for 3h at 25 °C. Cells were harvested at 6000xg for 15 min at 4 °C. Pellets were washed with ice-cold 50mM EDTA and centrifuged at 7000 rpm for 15 min at 4 °C. Washed pellets were saved at −80 °C until use.

#### Oligonucleotide site directed mutagenesis

Point mutation of aspartic acid residue D46 was generated by two step overlap PCR mutagenesis using primer pairs (KR38/KR41) and (KR40/KR39) with Phusion polymerase. A second PCR was performed with primers (KR38/KR39) using primary PCR products as a template and the subsequent PCR products were digested as above and ligated into pBAD24 to generate an N-terminal His6 tagged D46N YegK(PphC) fusion protein. Plasmid cloning was subsequently verified by restriction enzyme digest and DNA sequencing (Operon).

#### Purification of recombinant PphC

Frozen pellets were suspended in lysis buffer (20mM Tris pH 8.0, 250mM NaCl, 30mM imidazole, 10mM β-mercaptoethanol, 0.2% Triton-X 100, 10mg/ml lysozyme, 1mM PMSF) and incubated on ice for 30 min. Initial lysis was carried out by four cycles of freeze/ thaw in dry ice/ethanol bath and 37 ºC. Lysate was passed through a 22-gauge needle and added to pre-chilled 2mL screw cap tubes with 0.1mm silica beads. Cells were lysed using a FastPrep-24 5G instrument (MP Biomedical) using 2 rounds of 6.5m/s intensity for 40 secs with 4 min incubation on ice between rounds. Lysates were cleared by centrifugation at 20,000xg for 30 mins at 4 ºC. Cleared lysates were incubated in Pierce 5ml columns with Ni-NTA Agarose beads (Qiagen) at 4 ºC for 1hr. Lysate was allowed to flow through and beads were washed 10 column volumes of wash buffer (20mM Tris pH 8.0, 250mM NaCl, 30mM imidazole, 10mM β-mercaptoethanol). His6-tagged protein was eluted using increasing concentrations of imidazole (100–500mM) in 20mM Tris pH 8.0, 250mM NaCl. Elution fractions were analysed on a 10% SDS-PAGE gel and fractions containing protein were pooled and dialyzed overnight at 4 ºC using Slide-A-Lyzer mini dialysis device 10K MWCO (Thermo Scientific) in phosphatase storage buffer (20mM Tris pH 8.0, 50mMNaCl, 1mM DTT, 1mM MnCl2). Dialyzed samples were concentrated in Amicon Ultra 10K centrifugal filters to 1 ml and then loaded onto a HiLoad 16/60 superdex 75 prep grade (GE Biosciences) gel filtration column. The column was preequilibrated and run with phosphatase storage buffer. PphC elutes at a retention volume of ~70ml. Fractions containing PphC were pooled, concentrated and assessed for purity using a 12% SDS PAGE gel. The catalytic variant was purified using the same protocol. Proteins were stored at −80 ºC.

#### Cloning and expression of YegI

The *yegI* gene was PCR amplified using Phusion polymerase (Thermo Scientific) from *E. coli* REL606 genomic DNA using primers (KR58/KR59). Sequence for N-terminal His6 tag was included in the primer. The PCR product was digested with NcoI/SphI and ligated into similarly digested pBAD24 plasmid backbone. Ligation products were transformed in DH5a cells and selected on LB/ampicillin plates. The resulting plasmid generated an N-terminal His6-tagged YegI fusion protein. Plasmid cloning was subsequently verified by restriction enzyme digest and DNA sequencing (Operon). For protein expression, plasmid was transformed in *E. coli* LOBSTR (Kerafast) cells and plated on LB ampicillin (100μg/ml) agar plates. Single colonies were inoculated into 3ml LB supplemented with ampicillin (100μg/ml) for overnight cultures. The next day, a dilution of 1:250 in 500ml LB was grown to an OD600 of 0.6–0.8. Recombinant protein was induced by addition of arabinose (0.2% w/v) for 3h at 18 ºC. Cells were harvested at 6000xg for 15 min at 4 ºC. Pellets were washed with ice-cold 50mM EDTA and centrifuged at 7000 rpm for 15 min at 4 ºC. Washed pellets were saved at −80 ºC until use.

#### Purification of recombinant YegI

Frozen pellets were suspended in lysis buffer (50mM Tris pH 8.0, 200mM NaCl, 10mM ß- mercaptoethanol, 1mM PMSF and 2% w/v sarkosyl) and incubated at room temperature for overnight lysis. Cells were subsequently lysed using sonication. Lysates were cleared by centrifugation at 15000xg for 30 min at 4 °C. Cleared lysates were incubated in Pierce 5ml columns with Ni^2+^-NTA Agarose beads (Qiagen) at 4 °C for 1h. Lysate was allowed to flow through and beads were washed 10 column volumes of wash buffer (50mM Tris pH 8.0, 200mM NaCl, 30mM imidazole, 10mM β-mercaptoethanol, 0.05% w/v sarkosyl). His tagged protein was eluted using 300mM imidazole in 50mM Tris pH 8.0, 200mM NaCl, 10mM ß-mercaptoethanol and 0.05% w/v sarkosyl. Elution Fractions were tested on 12% SDS PAGE gel and fractions containing protein were pooled and dialyzed overnight at 4 °C using Snakeskin dialysis tubing 10K MWCO (Thermo Scientific) in kinase storage buffer (20mM Tris pH 8.0, 125mM NaCl, 10% glycerol,1mM DTT). Dialyzed protein was assessed for purity using 12% SDS PAGE gel and stored at −80 °C.

#### Phosphatase assays

The phosphatase activity of PphC (YegK) was determined by hydrolysis of p-nitrophenol phosphate (pNPP) to p-nitrophenol using spectrophotometry. Assays were performed in triplicate in a 96 well plate. Each well contained 350nM purified PphC (WT or mutant) in assay buffer (10mM T ris pH 8.0, 5mM MnCl2). Reactions were initiated by addition of 5mM pNPP (NEB) and absorbance measurements were recorded every 10 min at 405nm for 120 min in a Tecan Infinite 200 plate reader. Amount of phosphate released is represented as nmol pNP/μg of protein and the amount of pNP was calculated using extinction coefficient of 18000 M^−1^cm^−1^.

Metal dependence was determined by incubating 350nM purified PphC in assay buffer containing either 5mM MgCl2, 5mM MnCl2, 5mM CaCl2,5mM ZnCl2, 5mM NiCl2 or no metal. Absorbance measurements were recorded as above. To determine optimal MnCl2 concentrations, reactions were carried out with 350nM purified WT PphC in assay buffer with different MnCl2 concentrations (0–10mM) and absorbance was measured at 405nm.

Sensitivity to phosphatase inhibitors was determined by measuring phosphatase activity of purified PphC in the presence of the following: sodium phosphate (Sigma), sodium fluoride (Sigma), okadaic acid (Cell Signaling), sodium orthovanadate (NEB) or EDTA (Macron Chemicals), sanguinarine chloride (Tocris), aurin tricarboxylic acid (Alfa aesar) and 5,5’ methylene disalycilic acid (Acros organics). Okadaic acid, sanguinarine chloride, aurin tricarboxylic acid and 5,5’ methylene disalycilic acid were diluted in DMSO to get a 1 mM stock. The remaining inhibitors were diluted to stock concentrations in sterile water pH 8. 150nM of purified WT PphC was incubated in assay buffer containing indicated concentrations of inhibitor or DMSO/water for 5 min. Reactions were initiated by addition of 5mM pNPP and absorbance was recorded every 5 min for 15 min at 30 ºC in a Tecan Infinite 200 plate reader.

The kinetic parameters of PphC were determined by varying the pNPP concentration (0.1–6.4mM) in a reaction with 350nM of purified wild type PphC from either REL606 and MG1655 in assay buffer. Hydrolysis was monitored every 5 min for 30 min in the linear range of the reaction. Initial reaction velocities for every substrate concentration. To determine Km and Vmax values, the data was fit to a Michaelis-Menten curve and Lineweaver-Burke plot was derived using Graphpad Prism 7 software.

Synthetic phosphopeptides. To assess substrate specificity, 200μM of serine [RRApSVA], threonine [KRpTIRR] or tyrosine [RRLIEDAEpYAARG] phosphopeptides (Millipore) were incubated with 350nM phosphatase in a 50μl reaction containing 20mM Tris pH 8.0, 5mM MnCl2 for 30 min at 37 ºC. Phosphatase reaction was stopped by addition of Biomol Green reagent (Enzo). Reaction was incubated at room temperature for 25 min to allow color development and absorbance at OD620 was measured. Phosphate standard (Enzo) was used to calculate amount of phosphate released.

ß-casein. Phosphorylated β-casein (Sigma) was dissolved in 50mM Tris pH 7.5, 150mM NaCl to a final concentration of 4mg/ml. For the phosphatase assay, 4μg of β-casein was incubated with 1.5μg WT or D46N PphC in 20μl of 50mM Tris pH 8.0, 10mM MnCl2 at 37 ºC for 1 h. Reactions were stopped with 3X SDS loading dye and samples were heat denatured at 95 ºC for 5 min. Samples were loaded on 10% SDS polyacrylamide gel containing 50μM Phos-tag acrylamide as per manufacturer’s instructions (Wako). Gels were run at constant voltage of 150V for 75 min at 4 ºC. Proteins were visualized by Coomassie staining.

Dephosphorylation of autophosphorylated YegI. Autophosphorylation of YegI kinase was performed by addition of 5μCi of [g-^32^P] ATP (Perkin Elmer) to 2μM of purified YegI in 12μl of reaction buffer containing 50mM Tris pH 7.5, 50mM KCl, 0.5mM DTT, 10mM MgCl2, 10mM MnCl2, 200μM ATP. Reactions were incubated at 37 ºC and PphC or PphC-D46N was added to a final concentration of 4μM. Reactions were stopped at 45 min using 3X Laemmli buffer and boiled for 5 min. Samples were resolved on a 12% SDS-PAGE gel and visualized by straining with Coomassie dye. Radioactive dried gel was exposed and visualized by autoradiography.

## Acknowledgements

We would like to thank present and former members of our lab for helpful discussions. We acknowledge Elizabeth Nagle for performing some of the initial characterization of PphC.

This work was supported by an HHMI International Student Fellowship to KR and by NIH grant GM114213 to JD.

## Author contributions

KR performed all of the experiments and KR and JD wrote the manuscript.

---

**Table 1.** Inhibitor effects on PphC activity.

| No. | Inhibitors (Concentration) | % Relative activity |
| --- | --- | --- |
| 1. | No inhibitor | 100 |
| 2. | Sodium phosphate (5mM) | 13 |
|  | Sodium phosphate (10mM) | 6 |
| 3 | Sodium fluoride (10mM) | 81 |
|  | Sodium fluoride (100mM) | 70 |
| 4 | Sodium orthovanadate (5μM) | 87 |
|  | Sodium orthovanadate (50μM) | 95 |
| 5 | Okadaic acid (0.1μM) | 91 |
|  | Okadaic acid (1μM) | 88 |
| 6 | EDTA (1mM) | 63 |
|  | EDTA (2mM) | 38 |
| 7 | Aurin tricarboxylic acid (25μM) | 82 |
|  | Aurin tricarboxylic acid (50μM) | 66 |
|  | Aurin tricarboxylic acid (100μM) | 52 |
| 8 | 5,5' Methylene disalicylic acid (25μM) | 100 |
|  | 5,5' Methylene disalicylic acid (100μM) | 87 |
| 9 | Sanguinarine chloride (50μM) | 98 |

**Table 1: Effect of inhibitors on PphC phosphatase activity**

Reactions were carried out at 30 ºC for 15 min in buffer containing 150nM phosphatase with 5mM pNPP in the presence or absence of inhibitors. Relative activity was calculated as a percentage of phosphatase activity in the presence of inhibitor versus activity in absence of inhibito

